## Supplementary material for "Repression of Viral Gene Expression and Replication by the Unfolded Protein Response Effector XBP1u": Supllemental Tables S1, S2, S3

### Supplemental Material

| Name | Sequence |
|---|---|
| MIEP | CCATTATTGGC**ACGT**ACATAAGGTCAATAGGGGTGAGTCATTGGGTTTTTCCAGCCAATTTAATTAA<br>AACGCCATGTACTTTCCCACCATTG**ACGT**CAATGGGCTATTGAAACTAATGCA**ACGT**GACCTTTAA<br>ACGGTACTTTCCCATAGCTGATTAATGGGAAAGTACCGTTCTCGAGCCAATAC**ACGT**CAATGGGAA<br>GTGAAAGGGCAGCCAAAACGTAACACCGCCCCGGTTTTCCCCTGGAAATTCCATATTGGCACGCA<br>TTCTATTGGCTGAGCTGCGTTCT**ACGT**GGG**TATAA**GAGGCGCGACCAGCGTCGGTACCGTCGCA<br>GTCTTCGGTC |
| 1-mut<br>(motif 1 mutated) | CCATTATTGGCCTAGACATAAGGTCAATAGGGGTGAGTCATTGGGTTTTTCCAGCCAATTTAATTAA<br>AACGCCATGTACTTTCCCACCATTG**ACGT**CAATGGGCTATTGAAACTAATGCA**ACGT**GACCTTTAA<br>ACGGTACTTTCCCATAGCTGATTAATGGGAAAGTACCGTTCTCGAGCCAATAC**ACGT**CAATGGGAA<br>GTGAAAGGGCAGCCAAAACGTAACACCGCCCCGGTTTTCCCCTGGAAATTCCATATTGGCACGCA<br>TTCTATTGGCTGAGCTGCGTTCT**ACGT**GGG**TATAA**GAGGCGCGACCAGCGTCGGTACCGTCGCA<br>GTCTTCGGTC |
| 2-mut<br>(motif 2 mutated) | CCATTATTGGC**ACGT**ACATAAGGTCAATAGGGGTGAGTCATTGGGTTTTTCCAGCCAATTTAATTAA<br>AACGCCATGTACTTTCCCACCATTGCTAGCAATGGGCTATTGAAACTAATGCA**ACGT**GACCTTTAA<br>ACGGTACTTTCCCATAGCTGATTAATGGGAAAGTACCGTTCTCGAGCCAATAC**ACGT**CAATGGGAA<br>GTGAAAGGGCAGCCAAAACGTAACACCGCCCCGGTTTTCCCCTGGAAATTCCATATTGGCACGCA<br>TTCTATTGGCTGAGCTGCGTTCT**ACGT**GGG**TATAA**GAGGCGCGACCAGCGTCGGTACCGTCGCA<br>GTCTTCGGTC |
| 3-mut<br>(motif 3 mutated) | CCATTATTGGC**ACGT**ACATAAGGTCAATAGGGGTGAGTCATTGGGTTTTTCCAGCCAATTTAATTAA<br>AACGCCATGTACTTTCCCACCATTG**ACGT**CAATGGGCTATTGAAACTAATGCACTAGGACCTTTAA<br>ACGGTACTTTCCCATAGCTGATTAATGGGAAAGTACCGTTCTCGAGCCAATAC**ACGT**CAATGGGAA<br>GTGAAAGGGCAGCCAAAACGTAACACCGCCCCGGTTTTCCCCTGGAAATTCCATATTGGCACGCA<br>TTCTATTGGCTGAGCTGCGTTCT**ACGT**GGG**TATAA**GAGGCGCGACCAGCGTCGGTACCGTCGCA<br>GTCTTCGGTC |
| 4-mut<br>(motif 4 mutated) | CCATTATTGGC**ACGT**ACATAAGGTCAATAGGGGTGAGTCATTGGGTTTTTCCAGCCAATTTAATTAA<br>AACGCCATGTACTTTCCCACCATTG**ACGT**CAATGGGCTATTGAAACTAATGCA**ACGT**GACCTTTAA<br>ACGGTACTTTCCCATAGCTGATTAATGGGAAAGTACCGTTCTCGAGCCAATACCTAGCAATGGGAA<br>GTGAAAGGGCAGCCAAAACGTAACACCGCCCCGGTTTTCCCCTGGAAATTCCATATTGGCACGCA<br>TTCTATTGGCTGAGCTGCGTTCT**ACGT**GGG**TATAA**GAGGCGCGACCAGCGTCGGTACCGTCGCA<br>GTCTTCGGTC |
| 5-mut<br>(motif 5 mutated) | CCATTATTGGC**ACGT**ACATAAGGTCAATAGGGGTGAGTCATTGGGTTTTTCCAGCCAATTTAATTAA<br>AACGCCATGTACTTTCCCACCATTG**ACGT**CAATGGGCTATTGAAACTAATGCA**ACGT**GACCTTTAA<br>ACGGTACTTTCCCATAGCTGATTAATGGGAAAGTACCGTTCTCGAGCCAATAC**ACGT**CAATGGGAA<br>GTGAAAGGGCAGCCAAAACGTAACACCGCCCCGGTTTTCCCCTGGAAATTCCATATTGGCACGCA<br>TTCTATTGGCTGAGCTGCGTTCTCTAGGGG**TATAA**GAGGCGCGACCAGCGTCGGTACCGTCGCA<br>GTCTTCGGTC |
| all-mut<br>(all motifs mutated) | CCATTATTGGCCTAGACATAAGGTCAATAGGGGTGAGTCATTGGGTTTTTCCAGCCAATTTAATTAA<br>AACGCCATGTACTTTCCCACCATTGCTAGCAATGGGCTATTGAAACTAATGCACTAGGACCTTTAA<br>ACGGTACTTTCCCATAGCTGATTAATGGGAAAGTACCGTTCTCGAGCCAATACCTAGCAATGGGAA<br>GTGAAAGGGCAGCCAAAACGTAACACCGCCCCGGTTTTCCCCTGGAAATTCCATATTGGCACGCA<br>TTCTATTGGCTGAGCTGCGTTCTCTAGGGG**TATAA**GAGGCGCGACCAGCGTCGGTACCGTCGCA<br>GTCTTCGGTC |
| 4-only (all motifs mutated<br>except motif 4) | CCATTATTGGCCTAGACATAAGGTCAATAGGGGTGAGTCATTGGGTTTTTCCAGCCAATTTAATTAA<br>AACGCCATGTACTTTCCCACCATTGCTAGCAATGGGCTATTGAAACTAATGCACTAGGACCTTTAA<br>ACGGTACTTTCCCATAGCTGATTAATGGGAAAGTACCGTTCTCGAGCCAATAC**ACGT**CAATGGGAA<br>GTGAAAGGGCAGCCAAAACGTAACACCGCCCCGGTTTTCCCCTGGAAATTCCATATTGGCACGCA<br>TTCTATTGGCTGAGCTGCGTTCTCTAGGGG**TATAA**GAGGCGCGACCAGCGTCGGTACCGTCGCA<br>GTCTTCGGTC |

**Table S1.** Wildtyp and mutated Major Immediate-Early Promoter (MIEP) sequences. Minimal XBP1 binding motifs (ACGT) are underlined (black), mutated motifs are shown in red. The TATA box is in shown bold.

| Name | Orientation | Sequence |
|---|---|---|
| IRE1_1 | fwd | GCTTGCATGCTGTTAGCAAG |
| | rev | CTTGCTAACAGCATGCAAGC |
| IRE1_2 | fwd | GCATGCTGTTAGCAAGAGGA |
| | rev | TCCTCTTGCTAACAGCATGC |
| IRE1_3 | fwd | GATGGCAGTCTGTACACACT |
| | rev | AGTGTGTACAGACTGCCATC |
| XBP1_1 | fwd | GCGTAGACGTTTCCTGGCTA |
| | rev | TAGCCAGGAAACGTCTACGC |
| XBP1_2 | fwd | GGCTATGGTGGTGGTGGCAG |
| | rev | CTGCCACCACCACCATAGCC |
| XBP1_3 | fwd | GGTGGCAGCGGCGCCGAGCG |
| | rev | CGCTCGGCGCCGCTGCCACC |
| TRAF2_1 | fwd | GGAGCCAGGGGAAGTCACA |
| | rev | TGTGACTTCCCCTGGCTCC |
| TRAF2_2 | fwd | GAGGTACTTGGCTTCTAAC |
| | rev | GTTAGAAGCCAAGTACCTC |
| TRAF2_3 | fwd | GGGCCTGGAAAGGCCTCCGC |
| | rev | GCGGAGGCCTTTCCAGGCCC |

**Table S2.** gRNAs used for CRISPR/Cas9 gene editing.

| Name | Orientation | Sequence |
|---|---|---|
| Motif 1 | fwd | TCCATTATTGGCACGTACATAAGGTCATCCATTATTGGCACGTACATAAGGTCATCC ATTATTGGCACGTACATAAGGTCA-biotin |
| | rev | TGACCTTATGTACGTGCCAATAATGGATGACCTTATGTACGTGCCAATAATGGATGA CCTTATGTACGTGCCAATAATGGA |
| Motif 2 | fwd | TTCCCACCATTGACGTCAATGGGCTATTTCCCACCATTGACGTCAATGGGCTATTTC CCACCATTGACGTCAATGGGCTAT-biotin |
| | rev | ATAGCCCATTGACGTCAATGGTGGGAAATAGCCCATTGACGTCAATGGTGGGAAAT AGCCCATTGACGTCAATGGTGGGAA |
| Motif 3 | fwd | GAAACTAATGCAACGTGACCTTTAAACGAAACTAATGCAACGTGACCTTTAAACGAA ACTAATGCAACGTGACCTTTAAAC-biotin |
| | rev | GTTTAAAGGTCACGTTGCATTAGTTTCGTTTAAAGGTCACGTTGCATTAGTTTCGTTT AAAGGTCACGTTGCATTAGTTTC |
| Motif 4 | fwd | TCGAGCCAATACACGTCAATGGGAAGTTCGAGCCAATACACGTCAATGGGAAGTTC GAGCCAATACACGTCAATGGGAAGT-biotin |
| | rev | ACTTCCCATTGACGTGTATTGGCTCGAACTTCCCATTGACGTGTATTGGCTCGAACT TCCCATTGACGTGTATTGGCTCGA |
| Motif 5 | fwd | GAGCTGCGTTCTACGTGGGTATAAGAGGAGCTGCGTTCTACGTGGGTATAAGAGGA GCTGCGTTCTACGTGGGTATAAGAG-biotin |
| | rev | CTCTTATACCCACGTAGAACGCAGCTCCTCTTATACCCACGTAGAACGCAGCTCCTC TTATACCCACGTAGAACGCAGCTC |
| ERdj4 | fwd | CAACAGTTTTCCACGTGCGCGTAGGGCCAACAGTTTTCCACGTGCGCGTAGGGCCA ACAGTTTTCCACGTGCGCGTAGGGC-biotin |
| | rev | GCCCTACGCGCACGTGGAAAACTGTTGGCCCTACGCGCACGTGGAAAACTGTTGG CCCTACGCGCACGTGGAAAACTGTTG |
| LT ori | fwd | TAATTAAGCCTCTTAAGCCTCAGAGGCCTCTCTCTTTTTTTTCCAGAGGCCTCGGAG GCTAGGAGCCCCAAGCCTCTGCC-biotin |
| | rev | GGCAGAGGCTTGGGGCTCCTAGCCTCCGAGGCCTCTGGAAAAAAAAGAGAGAGGC CTCTGAGGCTTAAGAGGCTTAATTA |

**Table S3.** Oligonucleotides used for the DNA-Protein Interaction (DPI) ELISA.
